# Low and facultative mycorrhization of ferns in a low-montane tropical rainforest in Ecuador

**DOI:** 10.1101/2024.12.19.629367

**Authors:** Jennifer Michel, Marcus Lehnert, Martin Nebel, Dietmar Quandt

## Abstract

Arbuscular mycorrhizal fungi (AMF) are amongst the most studied plant symbionts and regularly found in terrestrial plants. However, global estimates of AMF abundance amongst all land plants are difficult because i) the mycorrhizal status of many non-commercial, wild plant species is still unknown ii) numerous plant species engage in facultative symbiosis, meaning that they can, but do not always do, associate with mycorrhiza and iii) mycorrhizal status can vary amongst individuals of one same plant species at one location, as well as for different plant species within a given genus or family.

To gain new insights to the pristine distribution of the plant-AMF symbiosis, we investigated the mycorrhizal status of one of the oldest lineages of extant vascular plants, Polypodiophyta (aka ferns), in one of the hotspots of natural plant diversification, the tropical rainforest. Providing a new data set of AMF abundance for 79 fern species, we hypothesized that (1) AMF would be found in 60-80% of the studied plants and (2) plant species with AMF symbionts would be more abundant than non-mycorrhizal species.

Both hypotheses were rejected while unexpected observations were made: (1) AMF occurred in 33% of studied species, representing 56% of the studied fern families, (2) AMF colonisation was not correlated with species abundance, (3) a small but significant proportion of AMF-hosting ferns was epiphytic (7%) and (4) mycorrhization was inconsistent among different populations of the same species (facultative mycorrhization).

Together this empirical data supports recent reservations regarding global abundance of AMF, and further demonstrates that mycorrhization is not a taxonomic trait. In addition, the occurrence of AMF in epiphytic plants and no net benefits of AMF for plant abundance indicate that the mycorrhization observed in this study is on the commensalism, possibly parasitism, side of the symbiosis spectrum.

**Highlights:**

- Small fraction of fern species mycorrhized (33% species level, 56% of families)
- Mycorrhization of species can vary with location (facultative symbiosis)
- AMF colonisation does not increase plant species abundance
- AMF presence in epiphytic ferns suggests commensalism / parasitism

## Introduction

Arbuscular mycorrhizal fungi (AMF) are amongst the most studied plant symbionts (Smith & Read, 2010). The inter-kingdom interaction between these fungi and plants is thought to be long-standing, with records of arbuscular structures inside plant roots dating back to 400-Million-year-old fossils (Strullu-Derrien et al. 2014; Sportes et al., 2021). The Devonian period (415-360 Ma) was characterized by an explosion in the diversity of land plants (Pires & Dolan, 2012) and the survival of the AM fungi-plant-relationship from the Devonian to present day times suggests an evolutionary benefit for both partners. The general principle of the textbook example of a mutualistic symbiosis is that the plant transfers carbon (C) from its photosynthetic activity to the rhizosphere, where C is exchanged with fungi. The fungi in turn help the plant to obtain nutrients like phosphorous (P) or nitrogen (N) from the soil matrix, which the fungi can release from soil minerals through mechanical or enzymatic activity of its hyphae (Landeweert et al. 2001; Cheng et al. 2012; Finlay et al. 2020). However, symbiotic interactions are nuanced and not always mutually beneficial.

The word “symbiosis” derives from the Greek “symbiōsis”, a combination of “syn”, meaning “with” and “bios”, meaning “life”, together meaning “state of living together” (Encyclopædia Britannica, 2023). When both partners benefit from the interaction, the relationship is termed “mutualism”. When one organism lives off another at the other’s expense, it’s called “parasitism”. When one species obtains food or other benefits from its symbiont, without either harming or benefiting the latter, it’s called “commensalism”. The character of a symbiotic interaction is case and context dependant, often a trade-off between dependence and benefit (Johnson et al., 1997; Purin & Rillig, 2007; Ramírez-Flores et al., 2020; Savolainen & Kytöviita, 2022). For example, the plant-AMF symbiosis involves mutualism, commensalism, parasitism and everything in-between (Neuhauser & Fargione, 2004), but in popular literature the term “symbiosis” often refers to only the narrow fraction of the whole symbiosis spectrum that consists of mutualism (Neubauer et al., 2024). Similarly, Karst et al. (2023) highlight an overly positive citation bias in studies investigating common mycorrhizal networks in forests and in their critical revision of both agronomic and mycorrhizal literature, Ryan and Graham (2018) show that no consistent link exists between yield benefits and AMF colonisation. AMF studies are also heavily biased geographically, with 92% of AMF investigations being conducted in the Northern hemisphere (Bennett & Classen 2020) and with unknown mycorrhization status for still 99% of plant species (Albornoz et al. 2021). This likely lead to an overestimation of the global occurrence of the AMF-plant-symbiosis and deep insights to the evolutionary trajectory of AMF are limited as the interaction is seldom studied at the hotspots of natural plant diversification which are located in the Global South, particularly along the Andean-Amazonian foothills (Corlett 2016; Sabatini et al. 2022; Albornoz et al. 2021). For *Plantago* for example, it has been shown that AMF abundance decreases with phylogenetic branch length, with indication that the reduction in AMF colonization might be driven by energetic costs (Formenti et al. 2023). Moreover, early AMF studies focussed on histological quantification of abundance of mycorrhizal structures, but lacked links to functioning or activity of AMF, and often also had no negative controls, which doesn’t allow conclusions about the character of the symbiosis along the mutualism-parasitism continuum (Neuhauser & Fargione 2004; Dudhane et al. 2024). A further important consideration for an evolutionary understanding of the AM fungi-plant-relationship is the fact that all AMF are plant-obligate symbionts, but only few plants are mycotroph (Brundrett & Tedersoo, 2018). Therefore, occurrence and functional and ecological integrity of AMF remain controversial topics.

This study addresses the data gap of AMF abundance in the South Hemisphere by providing new empirical data quantifying the presence of AMF amongst all fern families registered in a previous biodiversity survey in a low-montane tropical rain forest in eastern Ecuador (Michel et al., 2023). Ferns are an evolutionary old division and amongst the first vascular land plants, who appeared around 400 million years ago in the Devonian (Berry, 2009). Later in the Cretaceous, a fern radiation occurred giving rise to most modern fern families (Bomfleur et al., 2014). Ferns then became abundant floral elements in both temperate and tropical forests (Linares-Palomino et al. 2009; Chater, 2021). Given their wide distribution and versatility, ferns are also recognised as good indicators of overall biodiversity (Pouteau et al. 2016; Da Silva et al. 2018). Given that ferns strive despite the shade underneath the rainforest canopy and the highly weathered nutrient-poor soils of the study region, they are the ideal candidates to investigate plant-AMF symbiosis in a natural hotspot of plant diversification (Richter and Babbar 1991; Jobbágy and Jackson 2001; Cardoso and Kuyper 2016; Moreno-Jiménez et al. 2022).

In this study, two specific hypotheses were tested:

**(H1):** AMF will be highly abundant amongst ferns (60-80%).

**(H2):** Species with regular AMF symbiosis will be more abundant at plot level.

## Materials & methods

### Study area

The study area is located in eastern Ecuador in a remote and undisturbed part of the western Amazonian rainforest on the northern extensions of the foothills of the Cordilliera de Cutucú. The area is situated east of the national road E45 between El Puyo and Macas, between the rivers Pastaza and Macuma heading towards the township of Macuma (S02º06.664’ W077º44.334’). The land is part of the territory of the Shuar community of Wisuí, who kindly granted us access. The region is influenced by the high-pressure area over the central Amazon basin throughout the entire year and receives between 2500 mm and 4000 mm rainfall per year at moderate temperatures around 20°C. All sampling sites are located below 1500 m elevation and the soils are mostly entisols and inceptisol of low pH with considerable variation in exchangeable cations (Table 1). The climatic conditions remain relatively stable during the year and provide plants with an almost continuous vegetation period, making it a recognised hotspot of biodiversity and a habitat very suitable for ferns (Hawkins et al. 2003; Richter et al. 2009; Michel et al., 2023).

**Table 1:** Geochemical characterisation for the six studied plots. Abbreviations: C= Carbon, N= Nitrogen, C:N =carbon to nitrogen ratio, Al = aluminium, Ca = calcium, Fe = iron, K = potassium, Mg = magnesium, Mn = manganese; Na = sodium

| PLOT | Latitude | Longitude | Elevation<br>(m) | Soil type | pH<sub>H2O</sub> | C<br>(%) | N<br>(%) | C:N | Al | Ca | K | Mg | Mn | Na | $\Sigma$<br>Exchangeable<br>cations |
| --- | --- | --- | --- | --- | --- | --- | --- | --- | --- | --- | --- | --- | --- | --- | --- |
| | | | | | | | | | $\mu\text{mol g}^{-1}$ dry weight soil | | | | | | |
| WIS 1 | S 02° 06.803' | W 077°44.582' | 665 | Enti/Inceptisol | 5.26 | 5.50 | 0.50 | 10.97 | 4.98 | 186.52 | 2.40 | 11.54 | 0.33 | 0.66 | 206.43 |
| WIS 2 | S 02° 06.845' | W 077°44.587' | 658 | Alfisol | 6.82 | 7.95 | 0.69 | 11.48 | 0.00 | 391.35 | 3.36 | 19.29 | 0.05 | 0.58 | 414.64 |
| WIS 3 | S 02° 06.364' | W 077°45.788' | 996.5 | Enti/Inceptisol | 4.36 | 5.13 | 0.44 | 11.57 | 47.80 | 11.11 | 0.67 | 1.04 | 0.44 | 0.29 | 61.35 |
| WIS 4 | S 02° 06.361' | W 077°45.788' | 1002.5 | Enti/Inceptisol | 4.02 | 7.32 | 0.55 | 13.10 | 60.66 | 1.17 | 0.53 | 0.59 | 0.07 | 0.26 | 63.29 |
| WIS 5 | S 02° 06.000' | W 077°44.543' | 679.5 | Enti/Inceptisol | 4.18 | 7.39 | 0.57 | 12.89 | 69.86 | 32.81 | 2.43 | 10.37 | 0.47 | 0.28 | 116.22 |
| WIS 6 | S 02° 05.882' | W 077°44.529' | 688 | Enti/Inceptisol | 4.26 | 7.99 | 0.66 | 12.04 | 80.10 | 30.31 | 1.30 | 14.97 | 0.71 | 0.10 | 127.49 |

### Biodiversity survey and root collection

Ferns and their roots were collected in six plots established on the lower slopes of the mountain ridges (658 m – 1002 m). Each plot measured 20 m × 20 m (400 m^2^), and an additional collection was made on the mountain top of El Torre (1370 m). All fern species occurring within a plot were documented photographically and their life form was recorded (terrestrial, semi-epiphytic, epiphytic). Where available, four replicate plants were collected of each species in each plot, and plant material was conserved for distribution to the different collaborating herbaria (Quito, Macas, Stuttgart, Bonn). For each plot, alpha diversity was determined (number of different species per plot), and for each species their abundance (number of individuals of a species per plot) and relative abundance (the number of individuals of a species per plot relative to the total number of individuals per plot) were determined. Root samples were collected from each specimen. If several specimens were taken for a single species in one plot, respective root samples were combined into one root sample for that species in that plot. All root samples were first carefully cleaned and subsequently stored in 70% ethanol in Eppendorf tubes. In total, n=97 root samples were collected for AMF quantification. Terrestrial and epiphytic ferns had similar sampling size (n=44 terrestrial, n=11 semi-epiphytic, n=42 epiphytic). In addition to the biodiversity survey and the root samples, a composite sample of approximately 150 g of the upper soil horizon was taken in each plot for soil biochemical analysis.

### Root sample preparation and mycorrhiza documentation

To investigate presence/absence of AMF structures in root samples, the protocol after Grace & Stribley (1991) was used. Therefore, root samples were first transferred from the ethanol into 10% KOH and left for clearing 24h at 60°C. Then, root material was rinsed with distilled water twice and acidified with 10% lactic acid. As a dye, Aniline Blue WS (Sigma Aldrich, Missouri, US) was prepared in 0.05% lactic acid by dissolving 0.125 g solid Aniline blue in 125 ml distilled water and 125 ml lactic acid (0.125g/250ml). Roots were analysed in a two-step process: First, aliquots of 10 cm total root length of each sample were exposed to the staining solution for 45 min at 60°C. After washing with distilled water, these samples were used for screening and the detection of fungal hyphae. For this purpose, roots were cut into 10 pieces of 1cm length each. The 1 cm fragments were then aligned on microscope slides and gently prepared as squeeze fractions with the coverslip. In a second step, another aliquot of each root sample was analysed in cross sections to verify the localisation of arbuscules within parenchymal cells. For this purpose, untreated root material of each specimen was taken out of the ethanol and sliced into at least ten thin crosscuts, which were placed upon microscope slides. The thin cuts were covered with 10% KOH for 60 min and after acidification with 90% lactic acid, the procedure finished with the application of the Aniline Blue dye. All preparation steps were incubated on a heating plate at 45°C and constantly covered with a cover slide. Liquids were exchanged by pipetting them to one side of the coverslip while holding paper to the other side using capillary action. Visual inspection of both lateral squeeze sections and crosscuts was performed with a binocular stereomicroscope (Leitz F193, Leica Microsystems, Wetzlar, Germany; magnification: 16x), a light microscope (Leitz Laborlux S, Leica Microsystems, Wetzlar, Germany; magnification: 40x – 400x) with an Olympus D700 camera (Olympus Corporation, Tokyo, Japan) and a confocal microscope and its corresponding camera and software (VHX– 500I, Keyence, Osaka, Japan; magnification: 250x – 2500x). Each sample was than classified as either “mycorrhizal” or “non-mycorrhizal”. Species that occurred in several plots were classified as “facultative mycorrhizal” if they were “mycorrhizal” in one plot, but “non-mycorrhizal” in another.

## Data analysis

To test whether AMF root colonisation differed between plant growth forms (terrestrial, semi-epiphytic or epiphytic), and if relative abundance of fern species was influenced by mycorrhization (mycorrhizal, non-mycorrhizal, facultative mycorrhizal) we used analysis of variance. To test whether fern α-diversity was determined by the interaction between soil cation exchange capacity and AMF root colonisation we used analysis of co-variance with α-diversity as dependent variable and soil cation exchange capacity and AMF colonisation as covariates. All statistical analysis was carried out using R 4.4.1 (R Core Team, 2024) with the additional packages *car* (Fox & Weisberg, 2019) and *multcomp* (Hothorn et al., 2008).

## Results

### AMF at family and species level

Arbuscular mycorrhiza fungi were found colonizing individuals of nine fern families (Fig.1). Except for the Marattiaceae (Marattiopsida), all families are encompassed by the class Polypodiales (PPG I, 2016). Among these groups, the affinities to mycorrhization and the extent of mycorrhizal colonization differed between the investigated individuals. Depending on the plot, the sampled individuals of the different families belonged to either different or same genera and/or species. For example, the highest rate of mycorrhization was found for Marattiaceae, where all sampled rootlets presented AMF colonisation. All three samples were representatives of the genus *Danaea*. Strong mycorrhization was also found for Thelypteridaceae (85% of individuals investigated: 6 out of 7 different species of the genera Thelypteris), Dennstaedtiaceae (75% of individuals investigated: 3 of 4 different genera) and for Pteridaceae (75% of individuals investigated: 2 species of *Adiantum*, one species investigated twice). Again, two individuals of the same species were studied in the family of *Polypodiacea*, where individuals of *Nephrolepis rivularis* were studied from two different lowland plots. Mycorrhization was found only among rootlets of epiphytic plants of WIS5, whereas terrestrial individuals derived from WIS6 showed no mycorrhizal colonization of their roots. The opposite was detected among the *Athyriaceae*. Here, rootlets of *Diplazium ambiguum* were tested from epiphytic and terrestrial plants and only the ground-dwelling fern was colonised by AMF. Alike, two other terrestrial species of *Diplazium* (*D. expansum, D. macrophyllum*) were positively tested for root mycorrhiza. In the family of Dryopteridaceae, the terrestrial species *Polybotrya fractiserialis* was sampled twice at different sites within one plot (WIS5) and root analysis revealed AMF association for only one. Further mycorrhization was found for *Polybotrya polybotryoides, Didymochlaena truncatula* and for *Tectaria incisa*. Among the Lomariopsidaceae a variety of epiphytic *Elaphoglossum* species was investigated, but only Elapholossum ensiforme presented AMF. Among the other genera tested within this family, only the roots of the terrestrial *Lomariopsis prieuriana* were colonised by mutualistic fungi. Ultimately, AM fungi could be observed at the Polypodiaceae. Both representatives here grew semi-epiphytically in the same plot (WIS2). Individuals of *Campyloneurum phyllitidis* and *Serpocaulon adnatum* both grew as low epiphytes on dead wood as well as they were located on the ground.

**Figure 1:**
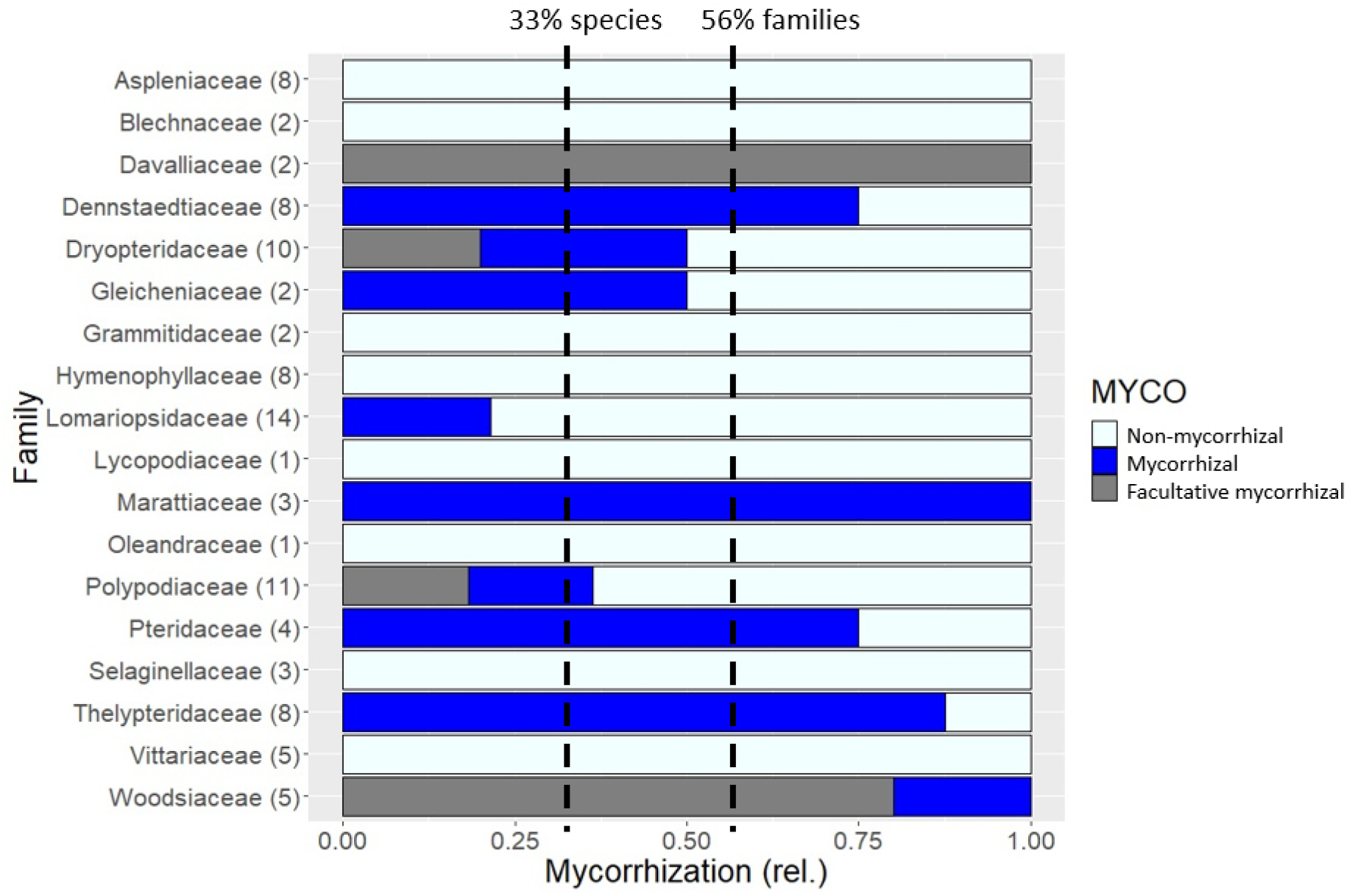
Abundance of AMF amongst 18 fern families of the Ecuadorian low-montane tropical rain forest. Number in brackets behind family name indicates number of species investigated within each family, total number of samples n=97. Dotted lines indicate average root colonisation with AMF across all investigated fern species (33%) and across all investigated fern families (56%) respectively.

### AMF occur in plants of all habitats but do not increase plant species abundance

Mycorrhizal colonisation was quantified for n=97 root samples from six survey plots in the area of Wisuí (WIS 1-6), comprising two mountain plots at an elevation above 900 m (WIS 3 and WIS 4) and four lowland plots located around 600 m above sea level (Table 1). Overall, 33% of all samples evidenced mycorrhizae. Terrestrial ferns species had significantly higher colonization levels (53.5%) than epiphytic ferns (7.3%). The plants of the intermediate habitats presented an intermediated level of mycorrhization, with 36.4% of the investigated semi-epiphytic fern individuals associated with symbiotic fungi (Fig.2A). There was no correlation between the mycorrhizal status of a fern species (mycorrhizal, non-mycorrhizal or facultative mycorrhizal) and the species’ relative abundance in a plot (Fig.2B). In an analysis of co-variance (Fig.2C), AMF remained unrelated to species abundance (p=0.22), but soil cation exchange capacity was correlated with species abundance (p=0.047).

**Figure 2:**
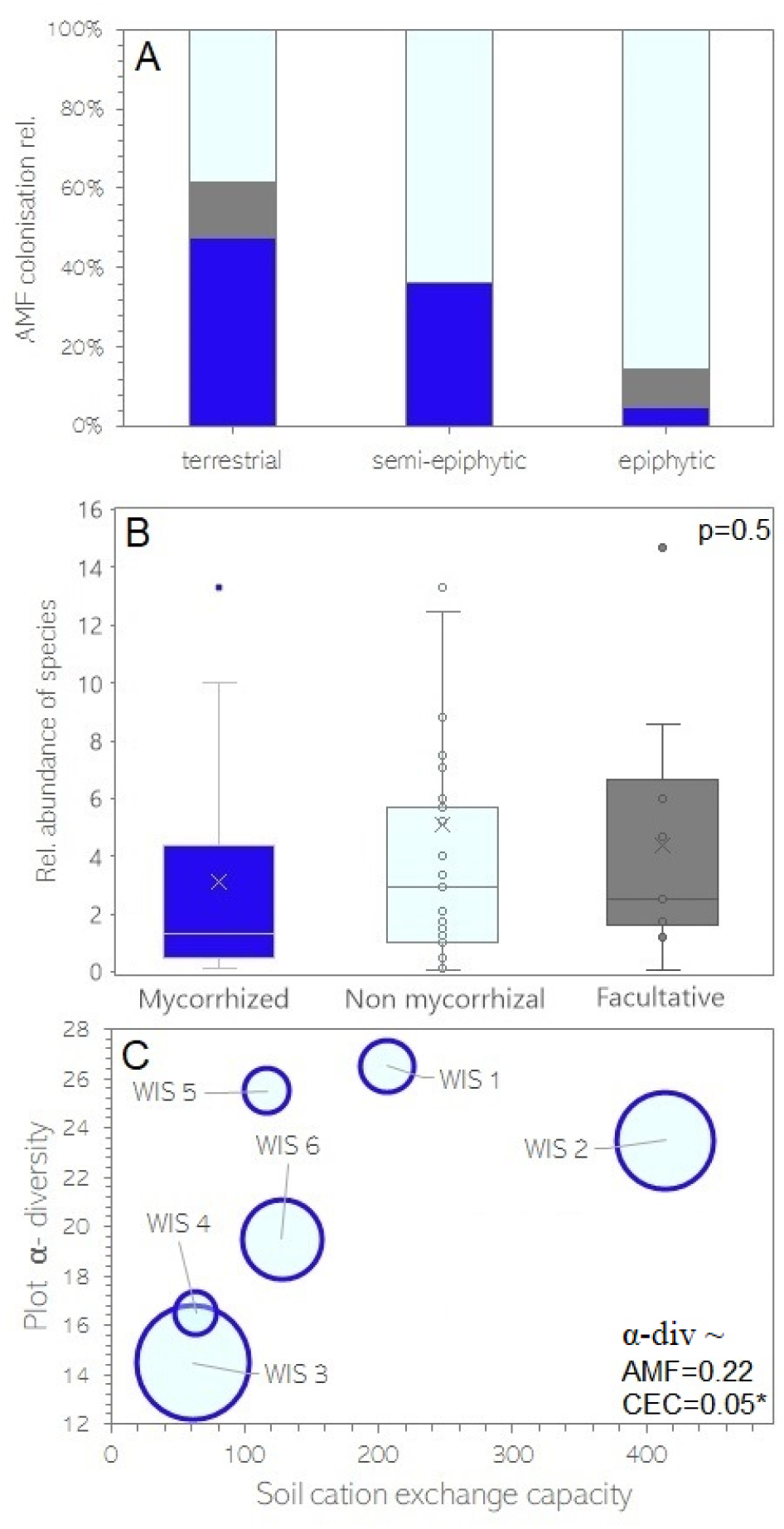
Relationship between (A) AMF colonisation and plant habitat (terrestrial, semi-epiphytic or epiphytic), (B) Relative abundance of fern species and their AMF status (mycorrhizal, non-mycorrhizal, facultative) with p-value of ANOVA and (C) Fern α-diversity at plot level, soil exchangeable cations (CEC) and degree of AMF-colonisation (indicated by the size of the circle) with p-value of analysis of co-variance with α-diversity as dependent variable and AMF and CEC as covariates.

## Discussion

### How abundant are AMF amongst land plants?

Most AMF research addresses economically important crops and angiosperms, while significantly less is known about AMF in other plant groups (Ganugi et al., 2019; Guillen et al., 2024). Albornoz et al. (2021) pointed out that abundance of AMF is globally overstated as the mycorrhizal status of 99% of plant species is still unknown and “mycorrhization” is easily overestimated as it is not a taxonomic trait, meaning that mycorrhization varies amongst species within plant genera and families, and even amongst individuals of the same plant species in the same location. AMF occur frequently in modern world angiosperms AMF, but in ancient groups like ferns they seem to be much less associated (Smith & Read, 2010). In this study we found 33% of fern species associated with AMF, representing 56% of the studied families. These observations are in line with a previous study that investigated AMF symbiosis in ferns in Southern Ecuador and found AMF in 29.10% of terrestrial species, equivalent to 48.49% of terrestrial samples (Lehnert et al., 2009) but lower compared to a global analysis reporting mycorrhization in 62% of fern species (Lehnert & Kessler, 2016). It has indeed been suggested that the abundance and reliance on AMF in ferns may have decreased during the evolutionary differentiation of fern genera as they occupied niches where they were more competitive without the fungal symbionts (Brundrett & Tedersoo, 2018; Watkins & Cardelús, 2012). Ferns might therefore rely more on other adaptive traits to compete, such as efficient spore dispersal, rapid growth in high-light conditions, and tolerance to less fertile soils. In addition, reduced dependence on fungal partners could have made ferns more resilient to environmental fluctuations or disruptions that might otherwise affect AMF communities and their dependant hosts.

### Facultative mycorrhization?

Facultative mycorrhization refers to plants that can form mycorrhizal associations with AMF but do not depend on them for survival. In this study five fern species were found to be facultatively mycorrhizal, namely *Diplazium ambiguum, D. hians, Microgramma thurnii, Nephrolepis rivularis* and *Polybotrya fractiserialis*. Facultative mycorrhization is actually very common among plants and has been previously described also for ferns (Guillen-Otero et al., 2024). For example, it has been observed that AMF colonisation increases with higher light intensity, likely related to an increased availability of photosynthetically fixed carbon compounds which are a main trademark in mutualistic symbiosis (Smith & Read, 2010; Guillen-Otero et al., 2024). In addition, changes in the community composition of AMF under varying light conditions were observed, which has also been reported for ferns grown on soil with varying nutrient levels and during seasonal changes from spring to winter (West et al., 2009; Zeng et al., 2024; Guillen-Otero et al., 2024). Finally, AMF have low taxonomic diversity and lack host specificity (Lee et al., 2013). Mycorrhization thus seems to be more driven by environmental factors than by host identity, which aligns with our finding that plant α-diversity is independent of AMF colonisation, but more driven by soil factors (Fig.2C). Accordingly, it has been observed that AMF association in ferns is independent of nitrogen and phosphorous fertilisation (Guillen et al., 2024). Future studies could address how common facultative mycorrhization is in angiosperms, as this type of symbiosis adds important nuance to global upscaling estimates of mycorrhization amongst plant species and families (Wang & Qiu, 2005). The environmental conditions under which abundance studies are conducted can influence recorded mycorrhizal colonization rates. For facultative mycorrhizal plants, favourable conditions might lead to higher observed associations, further complicating assessments of global AMF abundance. Understanding the ecological roles of AMF is crucial, as facultative associations may not reflect the true ecological significance of AMF in ecosystems.

### What do AMF do in the roots of epiphytic plants?

AMF associations in epiphytic plants are intriguing, as these plants live on other plants high up in the canopy rather than in the soil. The presence of AMF in the roots of these epiphytic plants could be an artefact, where future studies are needed to determine whether these fungi are dormant or parasitic. In other cases, canopy habitats can accumulate organic matter, creating a form of “canopy soil” (Nadkarni & Matelson, 1991). In these mini-ecosystems, AMF could facilitate nutrient transfer from decomposing materials, functioning similarly to the mutualistic symbiosis described for terrestrial ecosystems, but within the canopy structure. It is also possible that the AMF communities found in epiphytic ferns were in a dormant state (Bothe et al., 2010). Dormancy is common in AMF as it allows them to survive periods when environmental conditions like moisture and nutrient availability are not allowing active growth or symbiosis (Smith & Read, 2010). Finally, it might also well be possible that AMF in epiphytic plants are at the parasitism end of the symbiosis spectrum simply seeking for survival (Neuhauser & Fargione, 2004; Purin & Rillig, 2008). The presence of AMF in epiphyte roots also raises the question how the fungi arrived so high up in the canopy. AMF spores could reach epiphytic plants through wind dispersal, rain splash, or animal movement. Tree trunks and branches can act as conduits for spores to reach higher levels in the canopy, where they might form symbioses with epiphytes. Once in the canopy, AMF could sustain themselves on organic materials and humus, allowing them to persist and associate with epiphytic plants (Benzing, 2004; Snäll et al., 2005). Future studies could also try to answer the question what vertical distribution patterns AMF exhibit along trees and their epiphytes, ideally making use of molecular techniques.

### Are all AMF always actively involved in mutually beneficial symbiosis?

The observations of facultative mycorrhization and mycorrhization of epiphytic plants highlight the broad character of symbiosis in natural ecosystems, which spans a large spectrum from mutualism, over commensalism to parasitism, and certainly AMF associations with plants are not always mutually beneficial (Neuhauser & Fargione, 2004; Neubauer et al., 2024). Without further measurements, the sheer presence of AMF inside plant roots does not allow conclusions about whether the co-occurrence of these fungi inside plant structures has benefits for either of the two organisms. Hence, it is possible that AMF have been hitch-hiking plants since the Devonian, probably mostly as commensalists. To better understand the symbiosis, oservational data of root colonisation needs to be paired with plant nutrient uptake measurements and/or plant species abundance, with the relevant controls, to infer details about the character of the symbiosis. In their critical revision of both agronomic and mycorrhizal literature, Ryan and Graham (2018) show for example that no consistent link exists between yield benefits and AMF colonisation, and AMF are not required for optimal plant phosphorous nutrition. Sometimes AMF keep nutrients away from the plant, and sometimes reverse flow keeps water away from the plant (Querejeta et al., 2003; Hammer et al., 2011). In field studies, especially commercial AMF inoculum containing only a reduced number of fungal species seldom work (Säle et al., 2021). Competition between AMF species is also commonly reported (Engelmoer et al., 2014) and it has been shown that under elevated CO2 AMF can cause substantial losses of soil carbon (Cheng et al., 2012). AMF can also negatively impact their fellow soil organisms by keeping carbon away from them (Drigo et al. 2009). Another benefit attributed to AMF is their contribution to the formation and stability of soil aggregates (Barbosa et al. 2019), but this function can also be carried out by a plethora of different soil organisms (Camenzind et al. 2022), plant root mucilage (Shabtai et al. 2023), or by purely geochemical interactions between molecules in the soil matrix (McCully, 1999). Therewith, our results challenge the often stated essential purely positive role of AMF in ecosystems and call for more studies which i) investigate AMF abundance in undisturbed natural environments and ii) make clear links between presence and functionality of AMF, for example using stable isotope probing (Lekberg et al., 2013; Nuccio et al., 2022).

## Acknowledgments

We thank the Cotán family in Wisuí and the team of the herbarium in Quito for their warm welcome and safely guiding us through the jungle. We thank the team of the Natural History Museum in Stuttgart for their hospitality and advice during the study of AMF. The support of the DFG (grants LE1826/4 and QU153/7) and the BIOFAIR project (FNRS-R.8001.20) are greatly acknowledged. Research permits (No. 03-2012-Investigación-B-DPMS/MAE) were kindly issued by the Ministerio del Ambiente, Dirección Provincial del Morona-Santiago.

## References

Albornoz, F.E., Dixon, K.W. & Lambers, H. (2021). Revisiting mycorrhizal dogmas: Are mycorrhizas really functioning as they are widely believed to do?. Soil Ecol. Lett. 3, 73–82 10.1007/s42832-020-0070-2.

Barbosa M. V., de Fátima Pedroso D., Curi N., Carbone Carneiro M. A. (2019). Do different arbuscular mycorrhizal fungi affect the formation and stability of soil aggregates? (2019). Ciência e Agrotecnologia, 43:e003519. 10.1590/1413-7054201943003519.

Bennett AE, Classen AT. (2020). Climate change influences mycorrhizal fungal-plant interactions, but conclusions are limited by geographical study bias. Ecology 101(4):e02978. doi: 10.1002/ecy.2978.

Benzing DH (2004). Vascular epiphytes. In: Lowman MD, Rinker BH (eds) Forest canopies, 2nd edn. Elsevier, San Diego, pp 175–211. 10.1016/B978-012457553-0/50014-9.

Berry, Chris (2009). The Middle Devonian plant collections of Francois Stockman reconsidered. Geologica Belgica. 12 (1–2): 25–30.

Bomfleur, B.; McLoughlin, S.; Vajda, V. (2014). Fossilized Nuclei and Chromosomes Reveal 180 Million Years of Genomic Stasis in Royal Ferns. Science. 343 (6177): 1376–1377. Bibcode:2014Sci 343.1376B. doi:10.1126/science.1249884. PMID 24653037. S2CID 38248823.

Bothe H, Turnau K, Regvar M. (2010). The potential role of arbuscular mycorrhizal fungi in protecting endangered plants and habitats. Mycorrhiza (7):445–57. doi: 10.1007/s00572-010-0332-4.

Brundrett, M.C. and Tedersoo, L. (2018). Evolutionary history of mycorrhizal symbioses and global host plant diversity. New Phytol, 220: 1108–1115. 10.1111/nph.14976.

Camenzind, T., Mason-Jones, K., Mansour, I. et al. (2023). Formation of necromass-derived soil organic carbon determined by microbial death pathways. Nat. Geosci. 16, 115–122 10.1038/s41561-022-01100-3.

Cardoso IM and Kuyper TW (2006). Mycorrhizas and tropical soil fertility. Agriculture, Ecosystems and Environment 116, 72–84. https://www.sciencedirect.com/science/article/abs/pii/S0167880906001149.

Chater, C.C.C. (2021). Light in the darkness: How ferns flourished in the ancestral angiosperm forest. New Phytol, 230: 886–888. 10.1111/nph.17273.

Cheng, L., Booker, F.L., Tu, C., Burkey, K.O., Zhou, L., Shew, D.H., Rufty, T.W., Hu S?. (2012). Arbuscular Mycorrhizal Fungi Increase Organic Carbon Decomposition Under Elevated CO2. Science, 337, 6098: 1084–1087. https://www.science.org/doi/10.1126/science.1224304.

Christenhusz, M.J.M., Reveal, J.L., Farjon, A., Gardner, M.F., Mill, R.R. & Chase, M.W. (2011). A new classification and linear sequence of extant gymnosperms. Phytotaxa 19: 55–70. 10.11646/phytotaxa.19.1.3.

Corlett RT. (2016). Plant diversity in a changing world: Status, trends, and conservation needs. Plant Divers 38(1):10–16. doi:10.1016/j.pld.2016.01.001.

Cribari-Neto F, Zeileis A (2010). Beta regression in R. J Stat Softw. 10.18637/jss.v034.i02.

Da Silva VL, Mehltreter K and Schmitt JL (2018). Ferns as potential ecological indicators of edge effects in two types of Mexican forests. Ecological Indicators 93, 669–676. https://www.sciencedirect.com/science/article/abs/pii/S1470160×18303704?via%3Dihub.

Drigo, B., Pijl, A.S. Duyts et al. Kowalchuk, G.A. (2009). Shifting carbon flow from roots into associated microbial communities in response to elevated atmospheric CO2. PNAS 107 (24) 10938–10942 10.1073/pnas.0912421107.

Dudhane, M., Mahesh B. & Thomas, S. (2024). Advances in AMF Research: Isolation, Histochemical Staining, Enumeration, Morphological and Molecular Techniques. 10.1007/978-981-97-0296-1_2.

Encyclopaedia Britannica, The Editors of Encyclopaedia. “symbiosis”. Encyclopedia Britannica, 12 Sep. 2023, https://www.britannica.com/science/symbiosis. xAccessed 24 October 2023.

Engelmoer, D.J.P., Behm, J.E. and Toby Kiers, E. (2014). Intense competition between arbuscular mycorrhizal mutualists in an in vitro root microbiome negatively affects total fungal abundance. Mol Ecol, 23: 1584–1593 10.1111/mec.12451.

Finlay, R. D., Mahmood, S., Rosenstock, N., Bolou-Bi, E. B., Köhler, S. J., Fahad, Z., … & Lian, B. (2020). Reviews and syntheses: Biological weathering and its consequences at different spatial levels–from nanoscale to global scale. Biogeosciences, 17(6), 1507–1533.

Formenti, L, Iwanycki Ahlstrand, N. Hassemer, G, Glauser, G, van den Hoogen, J, Rønsted, N, van der Heijden, M, Crowther, TW, Rasmann, S (2023). Macroevolutionary decline in mycorrhizal colonization and chemical defense responsiveness to mycorrhization. iScience 26(5): 106632. 10.1016/j.isci.2023.106632.

Fox J, Weisberg S (2019). An R Companion to Applied Regression, Third edition. Sage, Thousand Oaks CA. https://www.john-fox.ca/Companion.

Ganugi, P., Masoni, A., Pietramellara, G., & Benedettelli, S. (2019). A review of studies from the last twenty years on plant–arbuscular mycorrhizal fungi associations and their uses for wheat crops. Agronomy, 9(12), 840.

Grace, C. & Stribley, D.P. (1991). A safer procedure for routine staining of vesicular-arbuscular mycorrhizal fungi, Mycological Research 95(10): 1160–1162. 10.1016/S0953-7562(09)80005-1.

Guillen T, Kessler M, Homeier J. (2024). Fern mycorrhizae do not respond to fertilization in a tropical montane forest. Plant Environ Interact. 5(2):e10139. doi: 10.1002/pei3.10139.

Guillen-Otero, T., Lee, SJ., Hertel, D. et al. (2024). Facultative mycorrhization in a fern (Struthiopteris spicant L. Weiss) is bound to light intensity. BMC Plant Biol 24, 103 10.1186/s12870-024-04782-6.

Hammer EC, Pallon J, Wallander H, Olsson PA. (2011). Tit for tat? A mycorrhizal fungus accumulates phosphorus under low plant carbon availability. FEMS Microbiology Ecology, 76(2): 236–244 10.1111/j.1574-6941.2011.01043.x.

Hawkins BA, Field R, Cornell HV, Currie DJ, Guégan J, Kaufman DM, Kerr JT, Mittelbach GG, Oberdorff T, O’Brien EM, Porter EE and Turner JRG (2003). Energy, water and broad-scale geographic patterns of species richness. Ecology 84, 3105–3117. 10.1890/03-8006.

Hothorn T, Bretz F, Westfall P (2008). Simultaneous Inference in General Parametric Models. Biometrical Journal, 50(3), 346–363.

Jobbágy EG and Jackson RB (2001). The distribution of soil nutrients with depth: global patterns and the imprint of plants. Biogeochemistry 53, 51–77. 10.1023/A:1010760720215.

Johnson, N.C., Graham, J.-H. and Smith, F.A. (1997). Functioning of mycorrhizal associations along the mutualism–parasitism continuum. New Phytol. 135: 575–585 10.1046/j.1469-8137.1997.00729.x.

Karst, J., Jones, M.D. & Hoeksema, J.D. (2023). Positive citation bias and overinterpreted results lead to misinformation on common mycorrhizal networks in forests. Nat Ecol Evo 10.1038/s41559-023-01986-1.

Landeweert, R, Hoffland, E, Finlay, RD, Kuyper, TW, van Breemen, N (2001). Linking plants to rocks: ectomycorrhizal fungi mobilize nutrients from minerals. Trends in Ecology & Evolution 16(5): 248–254. 10.1016/S0169-5347(01)02122-X.

Lee EH, Eo JK, Ka KH, Eom AH. (2013). Diversity of arbuscular mycorrhizal fungi and their roles in ecosystems. Mycobiology. (3):121–5. doi: 10.5941/MYCO.2013.41.3.121.

Lehnert, M., & Kessler, M. (2016). Mycorrhizal relationships in Lycophytes and ferns. Fern gazette, 20(3).

Lehnert, M., Kottke, I., Setaro, S., Pazmiño, L. F., Suárez, J. P., & Kessler, M. (2009). Mycorrhizal Associations in Ferns from Southern Ecuador. American Fern Journal, 99(4), 292–306. http://www.jstor.org/stable/25639842.

Lekberg, Y., Rosendahl, S., Michelsen, A., & Olsson, P. A. (2013). Seasonal carbon allocation to arbuscular mycorrhizal fungi assessed by microscopic examination, stable isotope probing and fatty acid analysis. Plant and Soil, 368, 547–555.

Linares-Palomino, R., Cardona, V., Hennig, E. I., Hensen, I., Hoffmann, D., Lendzion, J., … & Kessler, M. (2009). Non-woody life-form contribution to vascular plant species richness in a tropical American forest. Forest ecology: Recent advances in plant ecology, 87–99.

Martínez-García, L. & Pugnaire, FI (2011). Arbuscular mycorrhizal fungi host preference and site effects in two plant species in a semiarid environment. Applied Soil Ecology 48(3): 313–317. 10.1016/j.apsoil.2011.04.003.

McCully, M.E. (1999). Roots in soil: Unearthing the Complexities of Roots and Their Rhizospheres. Annu. Rev. Plant Physiol. Plant Mol. Biol. 50:695–718, 10.1146/annurev.arplant.50.1.695.

Michel J, Lehnert M, Quandt D. (2023). Elevation and cation exchange capacity determine diversity of ferns in a low-montane tropical rainforest in Ecuador. Journal of Tropical Ecology 39:e20. doi:10.1017/S0266467423000081.

Moreno-Jiménez E, Maestre FT, Flagmeier M, Guirado E, Berdugo M, Bastida F, Dacal M, Díaz-Martínez P, Ochoa-Hueso R, Plaza C, Rillig MC, Crowther TW and Delgado-Baquerizo M (2023). Soils in warmer and less developed countries have less micronutrients globally. Global Change Biology 29, 522–532. 10.1111/gcb.16478.

Nadkarni, N.M. and Matelson, T.J. (1991). Fine Litter Dynamics within the Tree Canopy of a Tropical Cloud Forest. Ecology, 72: 2071–2082. 10.2307/1941560.

Neubauer, A., Aros-Mualin, D., Mariscal, V. and Szövényi, P. (2024). Challenging the term symbiosis in plant–microbe associations to create an understanding across sciences. J. Integr. Plant Biol. 10.1111/jipb.13588.

Neuhauser C. & Fargione JE. (2004). A mutualism–parasitism continuum model and its application to plant–mycorrhizae interactions. Ecol. Mod. 177: 337–352 10.1016/j.ecolmodel.2004.02.010.

Nuccio, E. E., Blazewicz, S. J., Lafler, M., Campbell, A. N., Kakouridis, A., Kimbrel, J. A., … & Pett-Ridge, J. (2022). HT-SIP: a semi-automated stable isotope probing pipeline identifies cross-kingdom interactions in the hyphosphere of arbuscular mycorrhizal fungi. Microbiome, 10(1), 199.

Pires, N. D., & Dolan, L. (2012). Morphological evolution in land plants: new designs with old genes. Philosophical Transactions of the Royal Society B: Biological Sciences, 367(1588), 508–518.

Pouteau R, Meyer JY, Blanchard P, Nitta JH, TerorotuaMand Taputuarai R (2016). Fern species richness and abundance are indicators of climate change on high-elevation islands: evidence from an elevational gradient on Tahiti (French Polynesia). Climatic Change 138, 143–156. 10.1007/s10584-016-1734-x.

PPGI (2016). A community-derived classification for extant lycophytes and ferns. Journal of Systematics Evolution, 54: 563–603. 10.1111/jse.12229.

Purin S, Rillig MC. (2008). Parasitism of arbuscular mycorrhizal fungi: reviewing the evidence. FEMS Microbiol Lett. 279(1):8–14. doi: 10.1111/j.1574-6968.2007.01007.x.

Querejeta, J., Egerton-Warburton, L.M. & Allen, M.F. (2003). Direct nocturnal water transfer from oaks to their mycorrhizal symbionts during severe soil drying. Oecologia 134, 55–64 10.1007/s00442-002-1078-2.

R Core Team (2024). R: A Language and Environment for Statistical Computing. R Foundation for Statistical Computing, Vienna, Austria. https://www.R-project.org.

Ramírez-Flores MR, Perez-Limon S, Li M, Barrales-Gamez B, Albinsky D, Paszkowski U, Olalde-Portugal V, Sawers RJ. (2020). The genetic architecture of host response reveals the importance of arbuscular mycorrhizae to maize cultivation. Elife 9:e61701. doi: 10.7554/eLife.61701.

Richter M, Diertl KH, Emck P, Peters T and Beck E (2009). Reasons for an outstanding plant diversity in the tropical Andes of Southern Ecuador. Landscape Online 12. 10.3097/LO.200912.

Richter, D.D. & Babbar, L.I. (1991). Soil Diversity in the Tropics, Editor(s): M. Begon, A.H. Fitter, A. Macfadyen. Advances in Ecological Research, Academic Press (21): 315–389. 10.1016/S0065-2504(08)60100-2.

Ryan, M.H. and Graham, J.H. (2018). Little evidence that farmers should consider abundance or diversity of arbuscular mycorrhizal fungi when managing crops. New Phytol, 220: 1092–1107. 10.1111/nph.15308.

Sabatini, F.M., Jiménez-Alfaro, B., Jandt, U. et al. (2022). Global patterns of vascular plant alpha diversity. Nat Commun 13, 4683. 10.1038/s41467-022-32063-z.

Säle, V., Palenzuela, J., Azcón-Aguilar, C. et al. (2021). Ancient lineages of arbuscular mycorrhizal fungi provide little plant Benefit. Mycorrhiza 31, 559–576 10.1007/s00572-021-01042-5.

Savolainen, T. & Kytöviita, MM. (2022). Mycorrhizal symbiosis changes host nitrogen source use. Plant Soil 471, 643–654 10.1007/s11104-021-05257-5.

Shabtai, I.A., Wilhelm, R.C., Schweizer, S.A. et al. (2023). Calcium promotes persistent soil organic matter by altering microbial transformation of plant litter. Nat Commun 14, 6609 10.1038/s41467-023-42291-6.

Smith, S. E., & Read, D. J. (2010). Mycorrhizal symbiosis. Academic press.

Snäll T, Ehrlén J, Rydin H (2005). Colonization-extinction dynamics of an epiphyte metapopulation in a dynamic landscape. Ecology 86:106–115. 10.1890/04-0531.

Sportes, A., Hériché, M., Boussageon, R. et al. (2021). A historical perspective on mycorrhizal mutualism emphasizing arbuscular mycorrhizas and their emerging challenges. Mycorrhiza 31, 637–653 10.1007/s00572-021-01053-2.

Strullu-Derrien, C., Kenrick, P., Pressel, S., Duckett, J. G., Rioult, J. P., & Strullu, D. G. (2014). Fungal associations in Horneophyton ligneri from the Rhynie Chert (c. 407 million year old) closely resemble those in extant lower land plants: novel insights into ancestral plant–fungus symbioses. New Phytologist, 203(3), 964–979.

Wang, B., & Qiu, Y. L. (2006). Phylogenetic distribution and evolution of mycorrhizas in land plants. Mycorrhiza, 16, 299–363.

Watkins & Cardelús (2012). Ferns in an Angiosperm world: Cretaceous radiation into the epiphytic niche and diversification on the forest floor. International Journal of Plant Sciences 173(6). 10.1086/665974.

West B, Brandt J, Holstien K, Hill A, Hill M. (2009). Fern-associated arbuscular mycorrhizal fungi are represented by multiple Glomus spp.: do environmental factors influence partner identity? Mycorrhiza (5):295–304. doi: 10.1007/s00572-009-0234-5.

Zeng, K, Huang, D, Zhang, X, Liu, S, Huang, X, Xin, G (2024). Fern species and seasonal variation alter arbuscular mycorrhizal fungal colonization and co-occurrence patterns in the Heishiding Natural Reserve, South China, Applied Soil Ecology 193, 105172. 10.1016/j.apsoil.2023.105172.

